# Profiling Active Enzymes for Polysorbate Degradation in Biotherapeutics by Activity-Based Protein Profiling

**DOI:** 10.1101/2020.10.07.330076

**Authors:** Xuanwen Li, Divya Chandra, Simon Letarte, Gregory C. Adam, Jonathan Welch, Rong-Sheng Yang, Smaranda Bodea, Alex Dow, An Chi, Christopher A. Strulson, Douglas D. Richardson

**Affiliations:** Analytical Research & Development Mass Spectrometry, Merck & Co., Inc., 2000 Galloping Hill Road, Kenilworth, New Jersey 07033, United States; Biologics Process Research & Development, Merck & Co., Inc., 2000 Galloping Hill Road, Kenilworth, New Jersey 07033, United States; Quantitative Biosciences, Merck & Co., Inc., 770 Sumneytown Pike, West Point, PA 19486, United States; Biologics Analytical Research & Development, Merck & Co., Inc., 2000 Galloping Hill Road, Kenilworth, New Jersey 07033, United States; Chemical Biology, Merck & Co., Inc., 33 Avenue Louis Pasteur, Boston, MA 02115, United States

**Keywords:** Activity-Based Protein Profiling (ABPP), Biotherapeutics, Lipase, Polysorbate Degradation, Residual Host Cell Protein

## Abstract

Polysorbate is widely used to maintain stability of biotherapeutic proteins in formulation development. Degradation of polysorbate can lead to particle formation in drug products, which is a major quality concern and potential patient risk factor. Enzymatic activity from residual enzymes such as lipases and esterases can cause polysorbate degradation. Their high activity, even at low concentration, constitutes a major analytical challenge. In this study, we evaluated and optimized the activity-based protein profiling (ABPP) approach to identify active enzymes responsible for polysorbate degradation. Using chemical probes to enrich active serine hydrolases, more than 80 proteins were identified in harvested cell culture fluid (HCCF) from monoclonal antibodies (mAb) production. A total of 8 known lipases were identified by ABPP, while only 5 lipases were identified by a traditional abundance-based proteomics (TABP) approach. Interestingly, phospholipase B-like 2 (PLBL2), a well-known problematic HCP was not found to be active in process-intermediates from two different mAbs. In a proof-of-concept study, phospholipase A2 group VII (PLA2G7) and sialic acid acetylesterase (SIAE) were identified by ABPP as possible root causes of polysorbate-80 degradation. The established ABBP approach can fill the gap between lipase abundance and activity, which enables more meaningful polysorbate degradation investigations for biotherapeutic development.

## Introduction

Non-ionic surfactant polysorbate (PS) is an excipient in formulation solutions serving as a shear protectant to stabilize biotherapeutics, prevent agitation-induced aggregation, and minimize surface adsorption.^1-3^ Polysorbate-80 (PS-80) and polysorbate-20 (PS-20) are the most widely used PSs in the biopharma industry.^2-3^ The degradation of PS can result in turbidity and formation of sub-visible particles, impacting product quality.^1-2^ PS degradation can bring bioproduct critical quality attributes (CQA) out-of-specification by a few different mechanisms. PS degradation is an industry-wide challenge for process and formulation development in biotherapeutics.^1, 3-9^ The challenge increases with higher cell density and titer in bioprocess as well as higher drug concentrations in formulation development.^9-10^ There are multiple mechanisms for PS degradation which can be grouped into two main categories: oxidation and hydrolysis.^1, 3^ Enzymatic hydrolysis results in cleavage of an ester bond in PS by lipases, esterases or possibly other host cell proteins (HCPs) without known lipase activity. HCPs can co-purify with the therapeutic protein via specific or non-specific interactions.^11^ Several lipases, such as PLBL2, LPL and LPLA2, have been reported to degrade PS-80 or PS-20 in biotherapeutics.^4-7, 9^

Multiple analytical tools have been developed as part of an overall HCP control strategy.^11^ Total HCP testing via traditional enzyme-linked immunosorbent assay (ELISA) is currently considered the “gold standard”. ELISA provides one summed value for total HCP content. In some cases, specific antibodies were developed to known HCPs present in a sample.^12-13^ Alternatively, liquid chromatography-mass spectrometry (LC-MS) based proteomics approaches are evolving rapidly as an orthogonal assay for HCP characterization, since proteomics can identify and quantify practically any individual HCP with certain abundance levels.^14-18^ Absolute quantification of individual HCPs can be achieved by multiple reaction monitoring (MRM) or parallel reaction monitoring (PRM) targeted approaches with suitable internal standards.^6, 12^ However, current proteomics approaches have two main analytical challenges to overcome when investigating the cause of PS-80 degradation.

First, the detection limit of current proteomics strategies is limited by HCP abundance. Most proteomics-based approaches have detection limits at the sub-ppm level (HCP compared to drug substance, i.e. 6 orders of magnitude difference), which is limited by the dynamic range of the mass spectrometer.^17^ Some lipases, such as LPLA2, can significantly degrade PS while being present below sub-ppm concentration levels,^6^ thus going undetected by most proteomics platforms. To this end, research efforts have been focused on reducing the dynamic range of the samples to improve assay sensitivity, including limited native digestion,^16-17^ molecular weight cut-off^19^ or 2D-LC.^16, 18^ Second, enzyme abundance cannot tell the whole story of PS degradation. Different lipases/esterases may have different enzymatic activity and thus different impact on PS degradation. Determining enzyme activity is the key in understanding the root cause of the degradation since it does not always correlate with enzyme abundance.

Activity-based protein profiling (ABPP) is a chemical proteomic strategy for the characterization of functional states of enzymes in complex biological systems.^20-22^ ABPP offers an advantage over traditional abundance-based proteomics (TABP) techniques that rely on measuring abundance rather than the functional activity of target proteins. Synthetic chemical probes, consisting of a reactive group, a linker, and an enrichment tag are used to capture targeted active enzymes.^23^ The reactive group labels the active site of mechanistically related enzymes to form a covalent linkage. The covalently modified proteins can be detected or purified by the enrichment tag (e.g. biotin-avidin or alkyne-azide reaction) and identified by LC-MS. ABPP was first developed to investigate enzymatic activity of serine hydrolases,^24^ and cysteine proteases^25^ and has since been adapted for activity detection of more than a dozen enzyme classes, including proteases, kinases, phosphatases, glycosidases, and oxidoreductases.^26-28^ ABPP has enabled the study of enzyme activity in specific physiological and pathological processes on a proteome-wide scale. This approach was recently used for investigation of polysorbate degradation.^29^ By studying enzymes with probes that specifically label certain enzyme classes, previously uncharacterized enzymes have been assigned to a functional class.

Serine hydrolases are one of the largest known enzyme classes, comprising greater than 200 enzymes in humans.^21, 30^ It includes multiple lipases, esterases, thioesterases, amidases and peptidases,^30^ which are potentially harmful to biotherapeutics drug substances and formulation excipients. The defining characteristic of serine hydrolase superfamily members is the presence of a nucleophilic serine in the active site. This site can be covalently labelled with chemical probes for enrichment and subsequently identified by mass spectrometry.^20-22, 30^ In this study, we evaluated ABPP to identify active enzymes causing PS degradation in biologics.

## Materials and Methods

### Materials

Dithiothreitol (DTT) and high capacity streptavidin agarose were purchased from Pierce (Rockford, IL). Tris-HCI buffer and high-performance liquid chromatography (HPLC) grade solvents, DMSO, 10% SDS solution, acetonitrile (ACN) and Formic acid (FA) were purchased from Thermo Fisher Scientific (Waltham, MA). Monoclonal antibodies (mAbs) were obtained from the pipeline of Merck & Co., Inc., Kenilworth, NJ, USA under preclinical or clinical development. Universal Proteomics Standard 2 (UPS2) and Urea were purchased from Sigma (St. Louis, MO). Trypsin was purchased from Promega (Madison, WI). ActivX® Desthiobiotin-Fluorophosphonate (FP) was purchased from Thermo Fisher Scientific (Waltham, MA). FP-Biotin and thiazole urea probe were synthesized according to published reports.^24, 31^ Recombinant PLBL2, LPLA2, PLA2G7 and SIAE from Chinese hamster ovary (CHO) cells were expressed in CHO-3E7 cells and purified (>90% purity) by GenScript (Piscataway, NJ).

### Sample preparation for ABPP approach

Harvest cell culture fluid (HCCF) and products from the first ion-exchange column (IEXP) in downstream purification were used for ABPP testing. HCCF from CHO cells from fed-batch production of mAbs was diluted with 50 mM Tris (pH 8) to 2 mg/mL. IEXP samples were diluted to 10 mg/mL. Chemical probes were dissolved in dimethyl sulfoxide (DMSO) to make 0.1 mM stock solution. For each HCCF or IEXP sample (500 μL), 20 μL of chemical probe solution or control DMSO was added to make a final mixture concentration of 2 μM. All samples were incubated at room temperature for 2 hours with constant mixing on a rotator. After reaction, each sample was added 1 mL of ice-cold methanol/acetone (50:50) on ice for 30 minutes to remove unreacted probes. The precipitated proteins were collected via centrifugation at 20,000 *x g* for 15 minutes at 4 °C. The pellet was washed with 1 mL of ice-cold methanol/acetone solution once and dried under N_2_ gas. The remaining pellet was re-dissolved in 1 ml of sample suspension buffer (2M Urea, 1% SDS, 50 mM Tris, pH 8). For each sample, 60 μL of Pierce™ agarose beads were added before incubation at room temperature for 1 hours while constantly mixing on a rotator. The agarose beads were pelleted by centrifugation at 1000 *x g* for 1 minute at room temperature and washed with 500 μL of 2 M Urea/50 mM Tris 3 times and 50 mM Tris 2 times. The washed agarose beads were resuspended in 50 μL of 50 mM Tris-HCl. Trypsin (2 μg) was added and digested at 37 °C on a thermomixer at 500 rpm overnight. The supernatant was collected after centrifugation at 10,000 *x g* for 5 min at room temperature. The collected samples were reduced at 80 °C for 10 mins with 1 μl of 1 M DTT, and subsequently had 2 μL of 20% formic acid added before LC-MS analysis.

### LC-MS for ABPP analysis

The LC-MS was operated in nanoflow mode on an EASY-nLC 1200 System (Thermo, Bremen, Germany), coupled to a Q Exactive™ HF-X Hybrid Quadrupole-Orbitrap™ mass spectrometer (Thermo, Bremen, Germany). EASY-Spray C18 column was used (PepMap RSLC, ES803, 2 µm, 100 Å, 75 µm × 50 cm) with flow rate at 250 nL/min and column temperature at 40 °C. Mobile phase A was made of MS grade water with 0.1% FA, and Mobile phase B was made of MS grade 80% ACN with 0.1% FA. The gradient started with 2% B for 3 mins, and changed to 45% at 83 min, and increased to 70% at 98 mins. The column was washed with 100% B from 100 min to 110 min, followed by 2 cycles of zig-zag washing step from 2% B to 100% B to reduce carry-over peptides. The MS was run with data dependent analysis (DDA) with MS1 scan range from 350 to 2000 m/z and the top 10 most abundant ions for MS/MS fragmentation. The MS1 resolution was 60,000, AGC target 1e^6^ and maximum IT for 60 ms. The MS2 resolution was 15,000, AGC target 1e^5^ and maximum IT for 100 ms, isolation window at 1.4 m/z and NCE at 28. The dynamic exclusion was set for 10 s. The ESI source was run with sheath gas flow rate at 8, aux gas flow rate at 0, spray voltage at 3.8 kV, capillary temp at 320 °C, and Funnel RF level at 50.

### Sample preparation for TABP approach

For TABP approach, the sample preparation followed the native digestion method.^17^ Briefly, 1 mg of mAb process-intermediate samples from purification steps were incubated with trypsin (400:1, weight to weight) after pH adjust with 1M Tris-HCL for overnight digestion at 37 °C. To determine the limit of identification of the traditional proteomics workflow, UPS2, was spiked in a mAb drug substance before sample preparation. UPS2 is composed of 48 human proteins (6,000 to 83,000 Da) within a concentration range from 250 amol to 25 pmol. There are 8 proteins in each concentration group with 6 groups in total. After digestion, samples were denatured and reduced at 80 °C for 10 min with 2 µL of 50 mg/mL DTT. A large portion of undigested mAb was removed by centrifugation at 11,000 *x g* for 10 min. The supernatant was added 3 µL of 20% FA before LC-MS analysis.

### LC-MS for TABP analysis

LC-MS was performed on an ACQUITY UPLC H-Class system (Waters, Milford, MA) coupled with a Q Exactive™ HF-X Hybrid Quadrupole-Orbitrap™ mass spectrometer (Thermo, Bremen, Germany). The column was ACQUITY UPLC Peptide CSH C18 column (130 Å, 1.7 µm, 1 × 150 mm, Waters, Milford, MA) with flow rate at 50 µl/min and column temperature at 50 °C. Mobile phase A was MS grade water with 0.1% FA, and Mobile phase B was MS grade ACN with 0.1% FA. The gradient started with 1% B for 5 min, and changed to 5% at 6 min, and increased to 26% at 85 min. The column was washed with 90% B from 90 mins to 105 mins, followed by 2 cycles of zig-zag washing step from 5% B to 90% B to reduce carry-over peptides. The MS was run with DDA with MS1 scan range from 300 to 1800 m/z and top 20 for MS/MS fragmentation. The MS1 resolution was 60,000, AGC target 1e^6^ and maximum IT for 60 ms. The MS2 resolution was 15,000, AGC target 1e^5^ and maximum IT for 100 ms, isolation window at 1.4 m/z and NCE at 27. The dynamic exclusion was set for 20 s. The ESI source was run with sheath gas flow rate at 35, aux gas flow rate at 10, spray voltage at 3.8 kV, capillary temp at 275 °C, Funnel RF level at 35 and aux gas heater temp at 100 °C.

### Proteomics Identification and Data Analysis

MS raw data was searched against the internal CHO K1 fasta database from Merck & Co., Inc., Kenilworth, NJ, USA customized with mAb and spiked-in recombinant protein sequences using Proteome Discoverer 2.2. The precursor mass tolerance was set at 15 ppm and fragment mass tolerance at 0.02 Da. The dynamic modification was set for Met oxidation and maximum 3 modifications. The target FDR for peptide identification was 0.01 and protein identification filter required at least 2 unique peptides. The summed MS1 peak area from all identified peptides were used for estimation of protein relative abundance. In the ABPP approach, proteins with at least 4-fold enrichment compared to those from samples incubated with only DMSO were considered potential active serine hydrolases. The enriched proteins were searched again Ingenuity pathway analysis (QIAGEN, Redwood City, CA, USA) for their primary subcellular location and enriched molecular functions. Heatmap visualization was generated with R scripts.

### Lipase Activity Determination by PS-80 Force Degradation

The impact of lipase/esterase activity on PS-80 degradation was determined via an accelerated PS-80 incubation study. Briefly, PS-80 (at a concentration of 0.025%, w/v) was spiked into process intermediates (IEXP samples) or recombinant enzyme samples. The samples were then incubated at 37 °C and 70% humidity for up to 14 days. Two hundreds microliter aliquots of samples were taken at different time points and frozen at −80C. PS-80 concentration was determined by HPLC-CAD (Corona Charged Aerosol Detector, Thermo, Bremen, Germany) with Oasis Max column (2.1×20 mm, 30 μm, Waters, Milford, MA). Mobile phase A was 0.5% (v/v) acetic acid in water. Mobile phase B was 0.5% acetic acid in isopropyl alcohol. PS-80 was eluted with 20% to 70% mobile phase B from 3 to 11 mins. Standard curves with 25% to 150% of target concentration was run before and after testing samples to make standard curves for PS-80 concentration determination.

### Characterization of PS-80 Degradation by RP-UHPLC-ESI-HRMS

PS-80 purity was assessed with a Waters Acquity UPLC Classic coupled to a time□of□flight (TOF) mass spectrometer (Xevo G2-XS, Waters, UK) that was equipped with an ESI source. An Acquity UPLC Protein BEH C4 column (2.1 × 100 mm, 300 Å, 1.7 μm, Waters, Milford, MA, U.S.A.) was used for reversed-phase chromatography. Initial conditions were set at 98% solvent A (0.1% formic acid in water) and 2% solvent B (0.1% formic acid in acetonitrile). Solvent B was increased to 25% at 1.0 min, to 35% at 6.0 min, to 75% at 8.0 min, to 80% at 12.0 min, and to 99% at 17 min and held for 2 min, followed by equilibration step of 2% B for 4 min. The flow rate was 0.3 mL/min and column temperature was set at 50°C. One microliter of sample was loaded.

For the mass spectrometer, the following parameters were set for the ion source: capillary voltage 3.0 kV, sampling cone voltage 10 V, extraction cone voltage 3 V, source temperature 130°C, desolvation temperature 450°C, and cone gas flow rate at 30 L/h, desolvation gas flow rate at 850 L/h. Two MS functions with collision energy of 10 V and 20 V were used for promoting “in-source” fragmentation. ESI full□scan mass spectra were recorded over the m/z range of 100−4000 in resolution mode.

## Results

### ABPP Method Development for Profiling Serine Hydrolase Activity in Cell Culture from Biologics Production

Serine hydrolases, including lipases and esterases, can hydrolyze esters bonds in natural substrates, such as phospholipids or triglycerides, and release free fatty acids. These enzymes have been reported to have activity against PS-80 or PS-20 because they contain the same carboxylic ester bonds as their natural substrates (Figure 1A-B).^7^ The active site serine can be regenerated for the next reaction cycle through water-induced saponification of the acyl-enzyme intermediate (Figure 1A).^30^ The nucleophilic serine can be labelled and enriched using active site affinity probes containing an electrophilic group such as a fluorophosphonate (FP) or a triazole urea that covalently modifies the nucleophilic serine (Figure 1C-D).^26^

**Figure 1.**
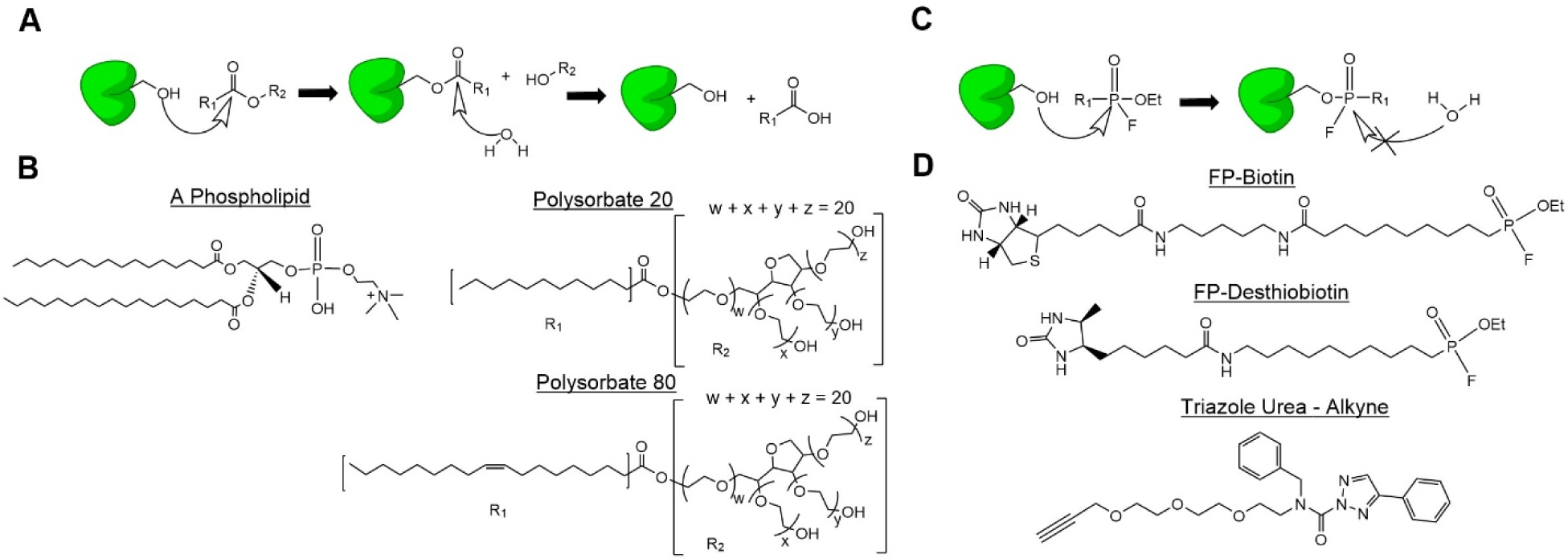
Mechanism of Actions of substrates and chemical probes reactions with serine hydrolases. (a) The generic mechanism of action for serine hydrolases. Taking note that the active-site serine can be regenerated by water induced saponification. (b) Structures of a generic phospholipid (a natural substrate), polysorbate-20 (PS-20) and polysorbate-80 (PS-80). All 3 compounds are hydrolyzed by serine hydrolases, with PS-20 and PS-80 being key excipients used in biotherapeutic formulations. (c) The generic mechanism of action for a chemical probe selective for serine hydrolases. (d) Structures of the probes used in this study. FP-Biotin, FP-Desthiobiotin and Triazole Urea are all active against serine hydrolases, whereas Triazole Urea is orthogonal and used as a control for non-specific binding in this study as it does not have the biotion group for subsequent enrichment with streptavidin agarose.

To establish the ABPP approach, two chemical probes (FP-Biotin and FP-Desthiobiotin) were evaluated using an HCCF sample of a mAb produced in CHO cells, referred to as mAb1 (Figure 2). The enrichment of serine hydrolases was compared to two controls: DMSO and an orthogonal triazole urea probe, used in this context as a broad-spectrum serine hydrolase inhibitor ^31^ (Figure 1D). As shown in Figure 2, the FP-Biotin probe shows stronger enrichment of serine hydrolases compared to the FP-Desthiobiotin probe. The enrichment capability of the two probes is most likely due to the higher affinity of biotin for streptavidin than desthiobiotin. The commercial FP-Desthiobiotin probe was used in a recent application of ABPP for PS-80 degradation.^29^ More active enzymes could probably have been identified if a more sensitive probe was used. More than 10 known lipases and esterases, were highly enriched with the FP-biotin probe. Some of those lipases have been reported to impact PS degradation, such as LPLA2 (PlA2G15, phospholipase A2, group XV)^6^, LPL (lipoprotein lipase)^5^ and the recently reported liver carboxylesterase^29^. Interestingly, another well reported and problematic lipase, PLBL2 (PLBD2, phospholipase B domain containing 2) ^10^, showed limited enrichment. Based on the enrichment performance of known serine hydrolases, FP-Biotin was selected for the rest of the ABPP studies.

**Figure 2.**
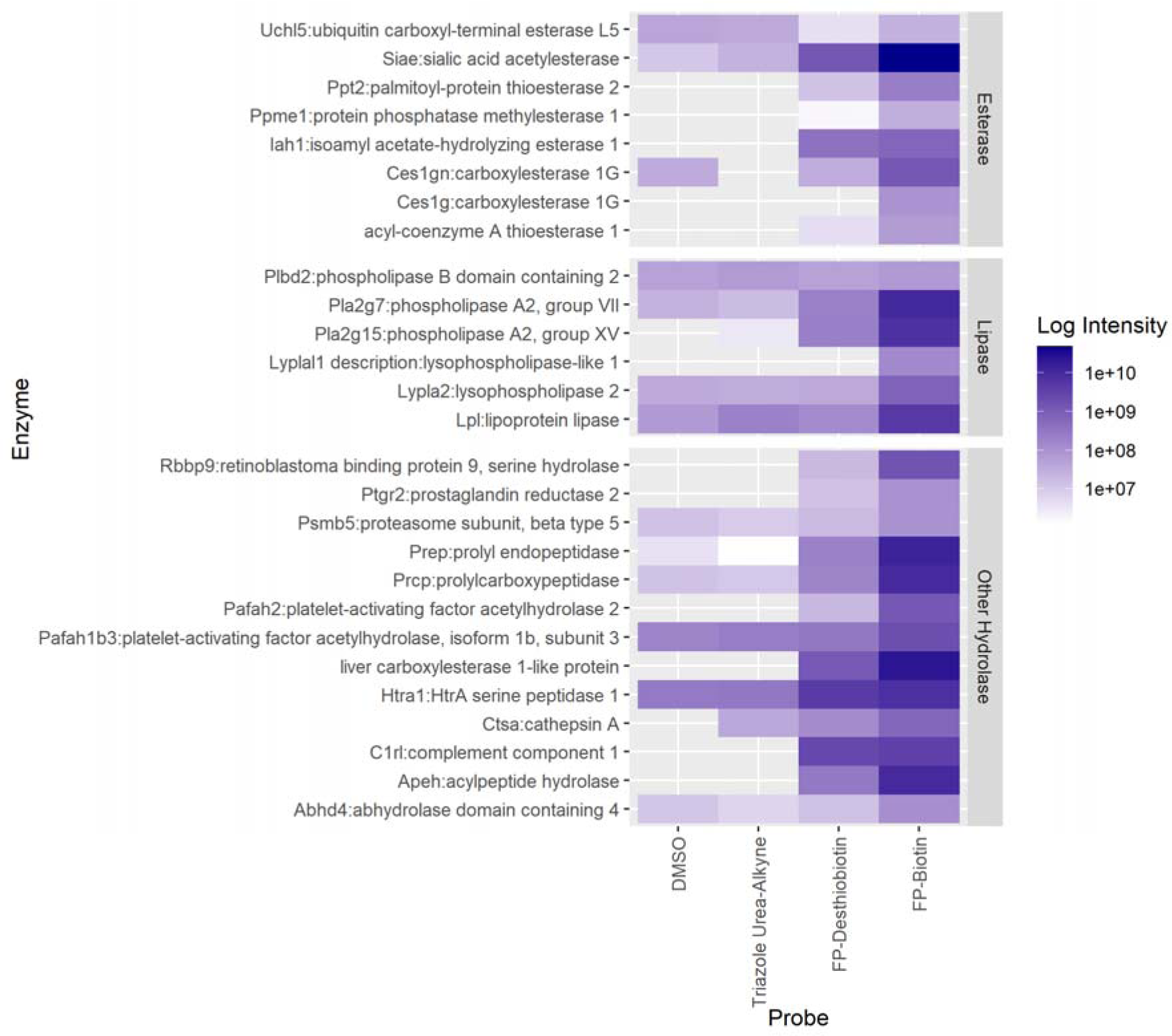
Evaluation of chemical probes using ABPP approach. Two positive probes (FP-Biotin and FP-Desthiobiotin) were evaluated using another probe (Triazole Urea) with different enrichment group and DMSO as controls. Several enriched hydrolases with functions as lipase and esterase were listed. The MS1 peak area was used for se-quantification comparison. Additionally, all enzymes are listed gene name followed by protein description.

To evaluate inter-day precision of the ABPP approach, HCCF samples from mAb1 were tested in two replicate experiments. As shown in Figure 3, comparison of the peak areas for commonly enriched proteins between the two preparations demonstrates assay reproducibility (R^2^>0.99).

**Figure 3.**
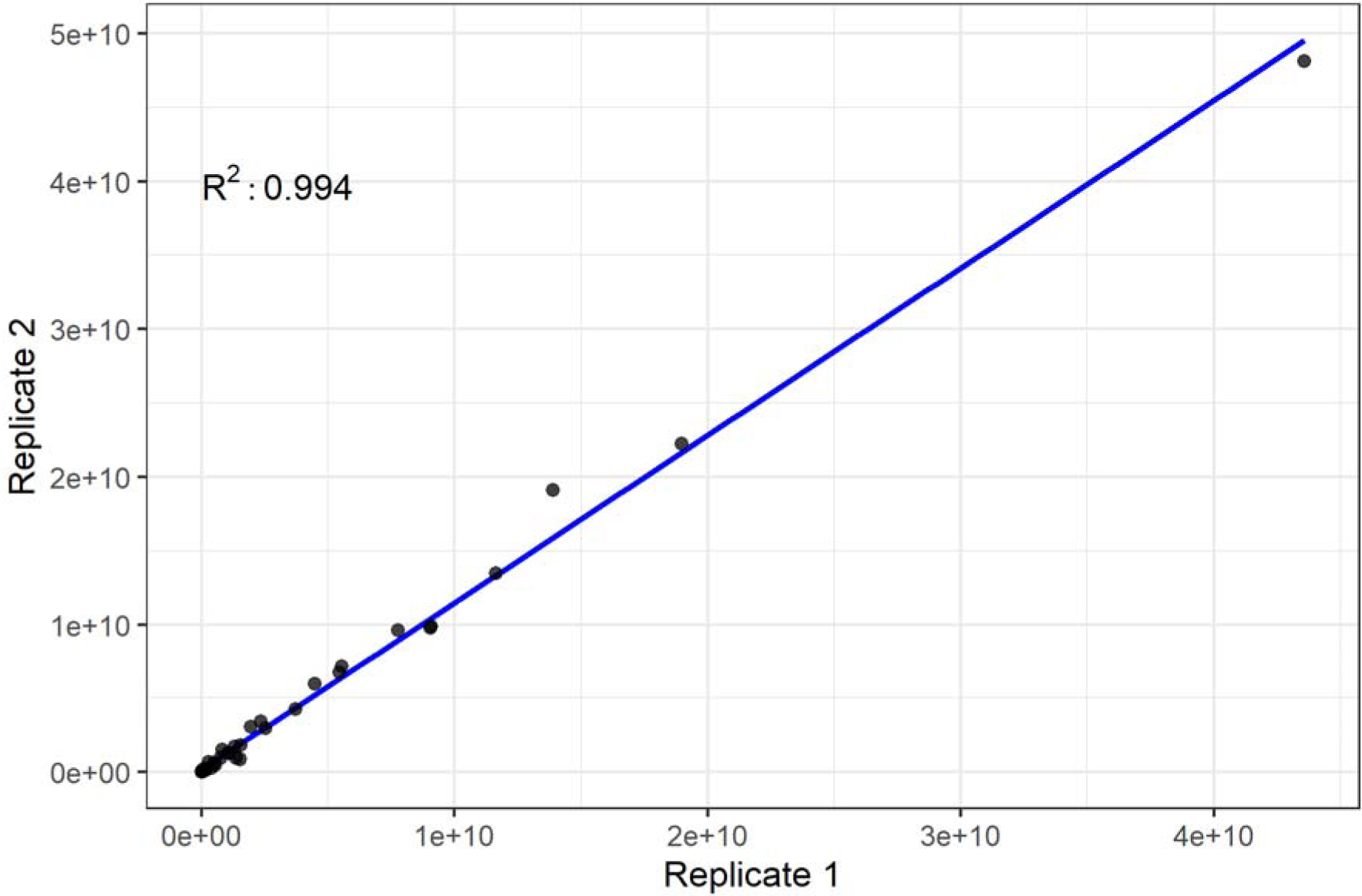
Interday precision of ABPP assay. HCCF sample from mAb1 was run with ABPP in two replicates. The peak area of those enriched proteins identified from the two experiments were compared to determine reproducibility of the technique.

### Advantages of ABPP Compared to TABP for Lipase Analysis

Keeping in mind the strengths of the ELISA for total HCP quantitation, we positioned the LC-MS based proteomics approach as an orthogonal assay for specific HCP characterization during process development. HCP identification for the shotgun-based TABP approach is biased on HCP abundance. To evaluate the identification limit of the established TABP approach, a UPS2 standard was spiked into a mAb drug substance before sample preparation and LC-MS analysis. All HCPs at more than 0.6 ppm were confidently identified with at least 2 unique peptides (Table 1 and Figure 4A). The average MS peak area from proteins of the same concentration group extracted by Proteome Discoverer 2.2 software correlates with the abundance of the spike-in protein in general (Table 1 and Figure 4B).

**Table 1.** Evaluation the detection limit of TABP approach using UPS2.

| UPS2 (fmol) | Range (ppm) | ID ($\geq 2$ unique peptides) * | Average MS1 peak area |
| --- | --- | --- | --- |
| 25,000 | 213-1659 | 8/8 | 6.82E+09 |
| 2500 | 16-149 | 8/8 | 7.98E+08 |
| 250 | 2-15 | 8/8 | 6.36E+07 |
| 25 | 0.2-1.2 | 5/8 | 1.29E+07 |

\*A total of 48 proteins in UPS2 was spiked in the testing. The remaining proteins were not identified or only identified by single unique peptides.

**Figure 4.**
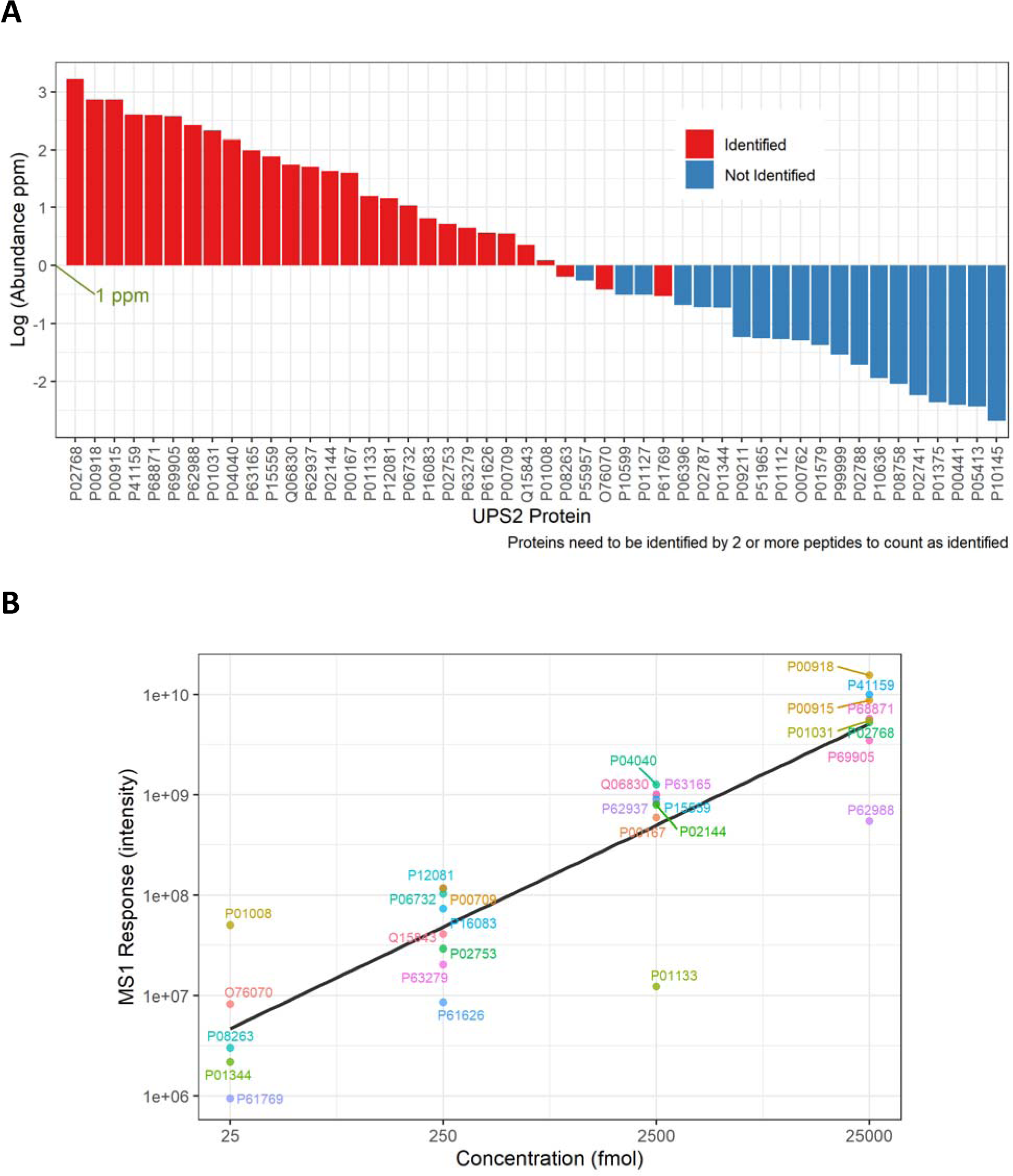
Spike-in UPS-2 to determine the sensitivity and semi-quantification of the TABP approach. (a) Spike-in proteins above 1 ppm were consistently identified with ≥ 2 unique peptides. (b) The average MS signal from MS1 peak area correlated very well with the spiked UPS2 protein concentration in general. The MS response varies among proteins from the same concentration range. Please note in (a) and (b) proteins are listed by UniProt ID.

To assess the advantage of ABPP compared to TABP for identification of low abundance HCP enzymes involved in PS degradation, the MS peak area of all the identified lipases from the two proteomic approaches in HCCF samples from two mAbs (mAb1 and mAb2) were compared. As shown in Table 2, ABPP resulted in better enrichment for most lipases compared to TABP. For example, the MS signal of LPLA2 (PLA2G15) from the ABPP was increased more than 150-fold compared with that from TABP. Studies have demonstrated that concentrations of LPLA2 below 1 ppm may have a significant impact on PS-80 degradation.^6, 32^ PLA2G7, a lipase from the same family as LPLA2, demonstrate the greatest enrichment (>350 fold) between the two methods. Several lipases were only identified utilizing ABPP including PLA1A (phospholipase A1 member A), LYPLA2 (lysophospholipase 2) and LYPLAL1 (lysophospholipase-like 1). The increase in MS signal may result from reduced complexity of the sample matrix as well as reduced ion suppression from the drug substance. Interestingly, some lipases were found to have a similar or even higher MS signal from TABP compared to ABPP, such as LIPA (lysosomal acid lipase A), LPL (lipoprotein lipase) and PLBL2 (phospholipase B domain containing 2). The similar MS signal intensity between the two approaches for LIPA and LPL may be due to their relative higher abundance levels in HCCF samples, and thus less ion suppression in MS. PLBL2 is the only lipase which has limited enrichment compared to DMSO control from both mAb HCCF samples. As a higher abundance protein in the sample, the presence of PLBL2 in the ABPP experiments at a lower concentration than in the TABP approach may be due to non-specific binding and interactions. This also suggests PLBL2 may not belong to the serine hydrolase family, or is not reactive with the probe molecule, or is present in the sample in an inactive form.

**Table 2.** The list of lipases identified from ABPP and TABP and their detected MS1 peak area from mAb1 and mAb2.

| Gene ID: Protein Description | mAb1 (ABPP) | mAb1 (TABP) | mAb2 (ABPP) | mAb2 (TABP) |
| --- | --- | --- | --- | --- |
| **PLA2G15**:phospholipase A2, group XV | 5.51E+09 | 3.65E+07 | 4.73E+09 | ND |
| **PLA2G7**:phospholipase A2, group VII | 9.57E+09 | 2.57E+07 | 8.91E+09 | 1.83E+07 |
| **PLALA**:phospholipase A1 member A | ND | ND | 3.26E+08 | ND |
| **LYPLA2**:lysophospholipase 2 | 1.45E+09 | ND | 6.58E+08 | ND |
| **LYPLAL1**:lysophospholipase-like 1 | 1.32E+08 | ND | 1.49E+08 | ND |
| **LPL**: lipoprotein lipase | 3.57E+09 | 3.54E+09 | 3.83E+08 | 8.74E+08 |
| **LIPA**:lysosomal acid lipase A | ND | 1.54E+08 | 2.19E+08 | 4.45E+08 |
| **PLBL2**:phospholipase B domain containing 2 | 5.26E+07 | 2.16E+09 | 2.27E+07 | 1.54E+08 |

ND: not detected

### Profiling Serine Hydrolases from two mAbs Bioprocesses

To understand the overall protein composition of proteins enriched by ABPP, HCCF samples from two different mAbs and different CHO cells were tested. Of all the 82 proteins enriched more than 4-fold compared to DMSO control by the FP probe from at least one of the mAb HCCF samples, 55 proteins (>67%) are annotated as hydrolases, and 27 of 82 proteins have unannotated functions as hydrolases. Most of the enriched proteins were annotated as cytoplasmic molecules for their primary subcellular locations from Ingenuity Pathway Analysis, a few proteins were annotated as extracellular and membrane proteins (Figure 5). The biological function annotations for the identified proteins were highly enriched for proteolysis (p value = 8.69E-10) and lipolysis (p value = 1.19E-8).

**Figure 5.**
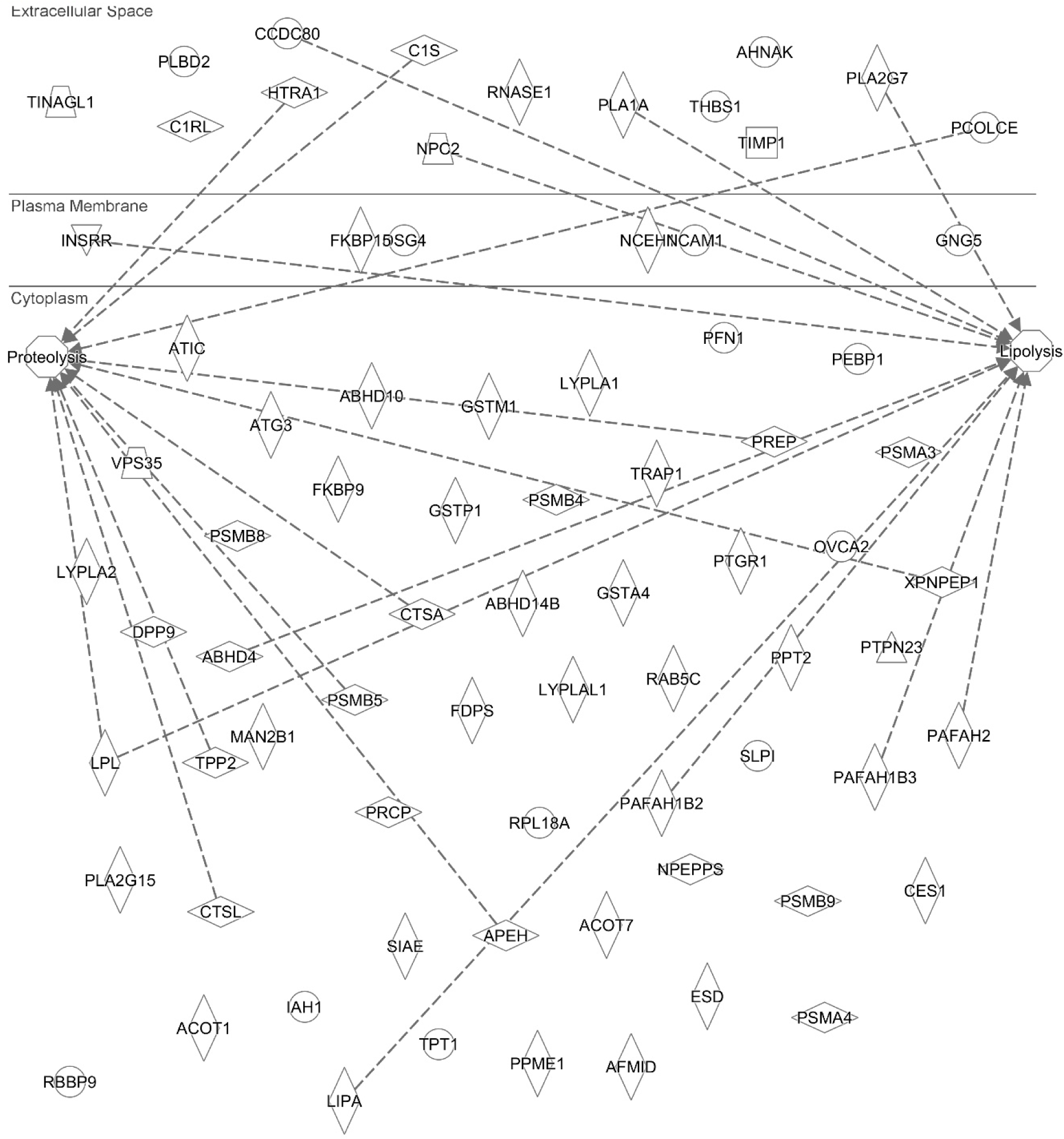
The subcellular locations and enriched functions of the identified proteins from ABPP by Ingenuity pathway analysis. Proteins are displayed by their gene ID.

In order to compare the serine hydrolase profile between different CHO cell lines, the enriched proteins from the ABPP assay were compared between mAb1 and mAb2. More than 36% of the enriched hydrolases are only identified from mAb1 or mAb2, or have a greater than 4-fold difference in enrichment in one or the other cell line. The remaining HCPs are within 4-fold in the HCCF samples of the two mAbs (Figure 6).

**Figure 6.**
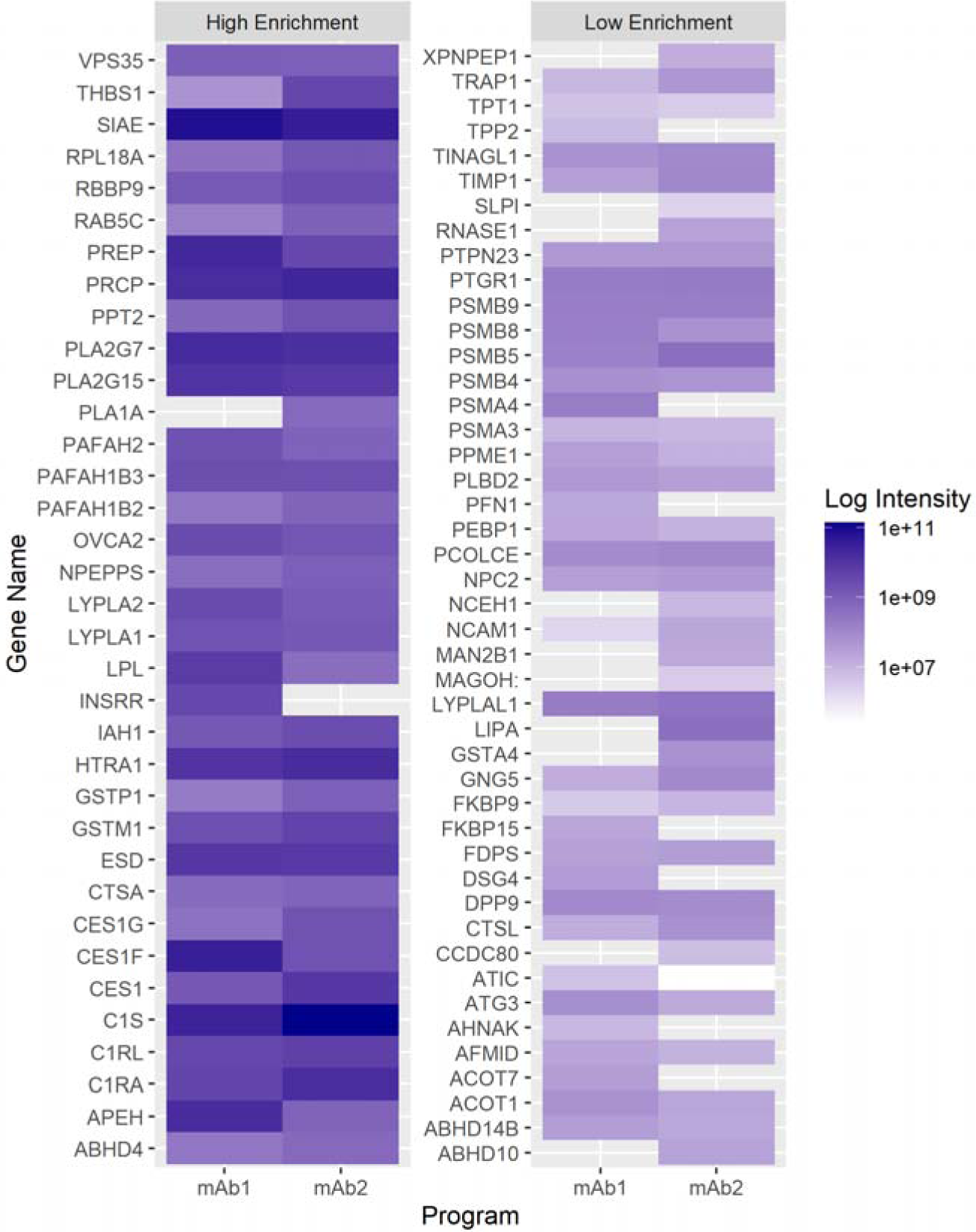
The comparison of serine hydrolases profile in HCCF from two mAbs by the ABPP. The MS1 peak area was used for semi-quantification comparison.

### ABPP for Root Cause Investigation of PS-80 Degradation

To demonstrate the application of ABPP for process characterization and root cause investigations during bioprocess development, the IEXP samples from mAb1 and mAb2 were analyzed by both TABP and ABPP approaches. IEXP samples are obtained after purification of HCCF via Protein A chromatography followed by an ion exchange chromatography polishing step. Thus, IEXP samples are enriched for the desired mAb and contain much less HCPs than HCCF samples, both in HCP numbers and in abundance. PS-80 incubation studies indicated there was significant degradation for mAb2 and minimal degradation for mAb1 compared to the placebo control that doesn’t contain drug substance (Table 3). The TABP method failed to identify any lipase contaminants in mAb2, but PLBL2 was identified in mAb1. The PLBL2 level in mAb1 was determined to be 64 ppm by an in-house PLBL2 ELISA assay.^12^ By contrast, ABPP did not identify any enriched enzymes from mAb1. This corroborates our previous result showing that PLBL2 is not enriched by the activity-based probe. Interestingly, two serine hydrolases, PLA2G7 (phospholipase A2, group VII) and SIAE (sialic acid acetylesterase), were identified in IEXP of mAb2 by ABPP.

**Table 3.** Comparison of identified lipase/esterase by ABPP and TABP from the PS-80 degradation study.

| **IEXP** | **LC-MS based Proteomics** |  | **%PS-80 degradation** |
| --- | --- | --- | --- |
|  | **TABP** | **ABPP** | **37° C/day 14** |
| **mAb1** | **PLBL2**<br>(64 ppm) | No lipase identified | 6.67 |
| **mAb2** | No lipase identified | **PLA2G7, SIAE** | 27.27 |

### Confirmation of PS-80 Degradation by Recombinant enzymes

To determine that the active lipases and esterase identified by ABPP may be involved in PS-80 degradation, four recombinant enzymes PLBL2, LPLA2, PLA2G7 and SIAE purified from CHO cells were spiked into mAb2 samples at 15 g/L. Herein, the mAb2 samples selected for the study were purified via an improved downstream purification process and thus expected to have even lesser HCPs than the mAb-2 IEXP samples where PLA2G7 and SIAE were identified via ABPP as described above. The rationale behind spiking in recombinant enzymes into a mAb sample, was to mimic the typical matrix obtained after downstream purification. As shown in Figure 7, LPLA2 with known activity against PS-80 shows time- and concentration-dependent PS-80 degradation. The newly identified lipase PLA2G7 had similar impact on PS-80 degradation as LPLA2. As consistent with the data from ABPP profiling as shown in Figure 2, PLBL2 had no activity on PS-80 degradation. Recombinant SIAE also had no impact on PS-80 degradation. This could be possibly due to low quality of recombinant SIAE with limited overall activity. PS-80 characterized by LC-MS (Figure 8) in the samples spiked with recombinant enzyme confirmed PS-80 degradation, which was consistent with results from analysis with HPLC-CAD. Mono-esters from PS-80 were the major species degraded by LPLA2 and PLA2G7 (Figure 8D and 8E).

**Figure 7.**
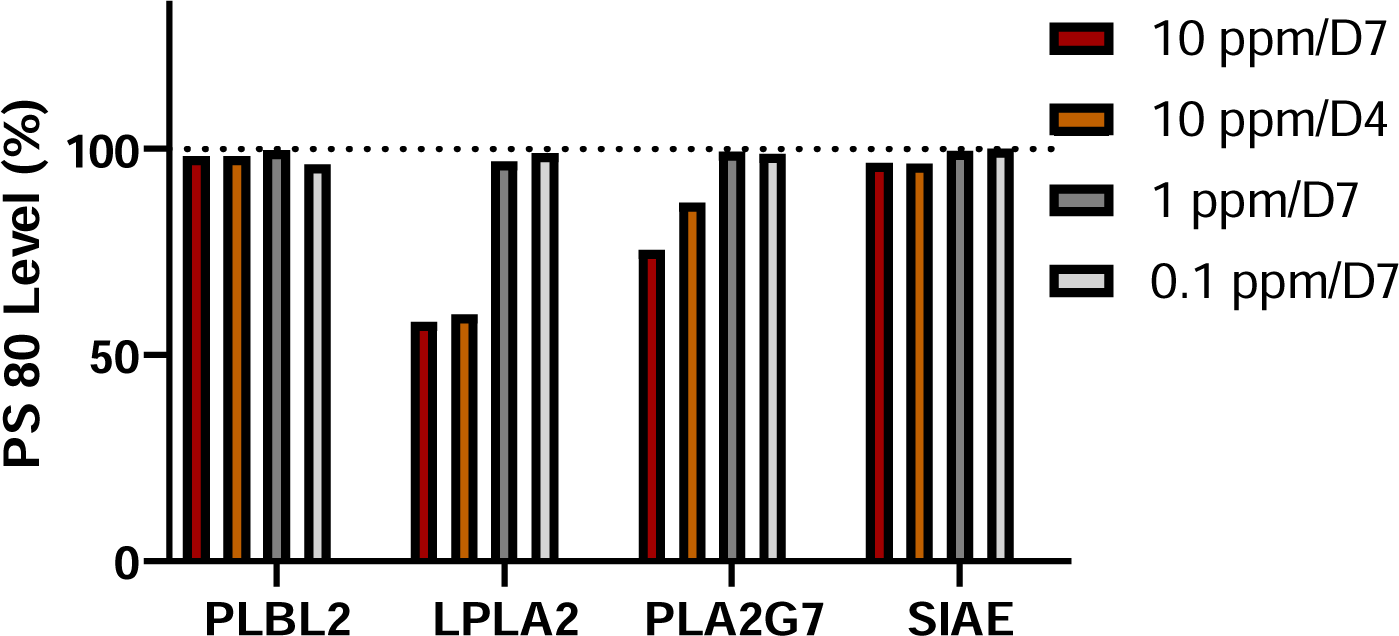
The impact of recombinant lipase or esterase on PS 80 degradation. Various concentrations (0.1 to 10 ppm) of recombinant enzyme was added in purififed mAb2 from an improved process and incubated with 0.02% PS 80. PS-80 levels measured by HPLC-CAD from day 4 (D4) and day 7 (D7) were compared to samples without spike-in enzymes.

**Figure 8.**
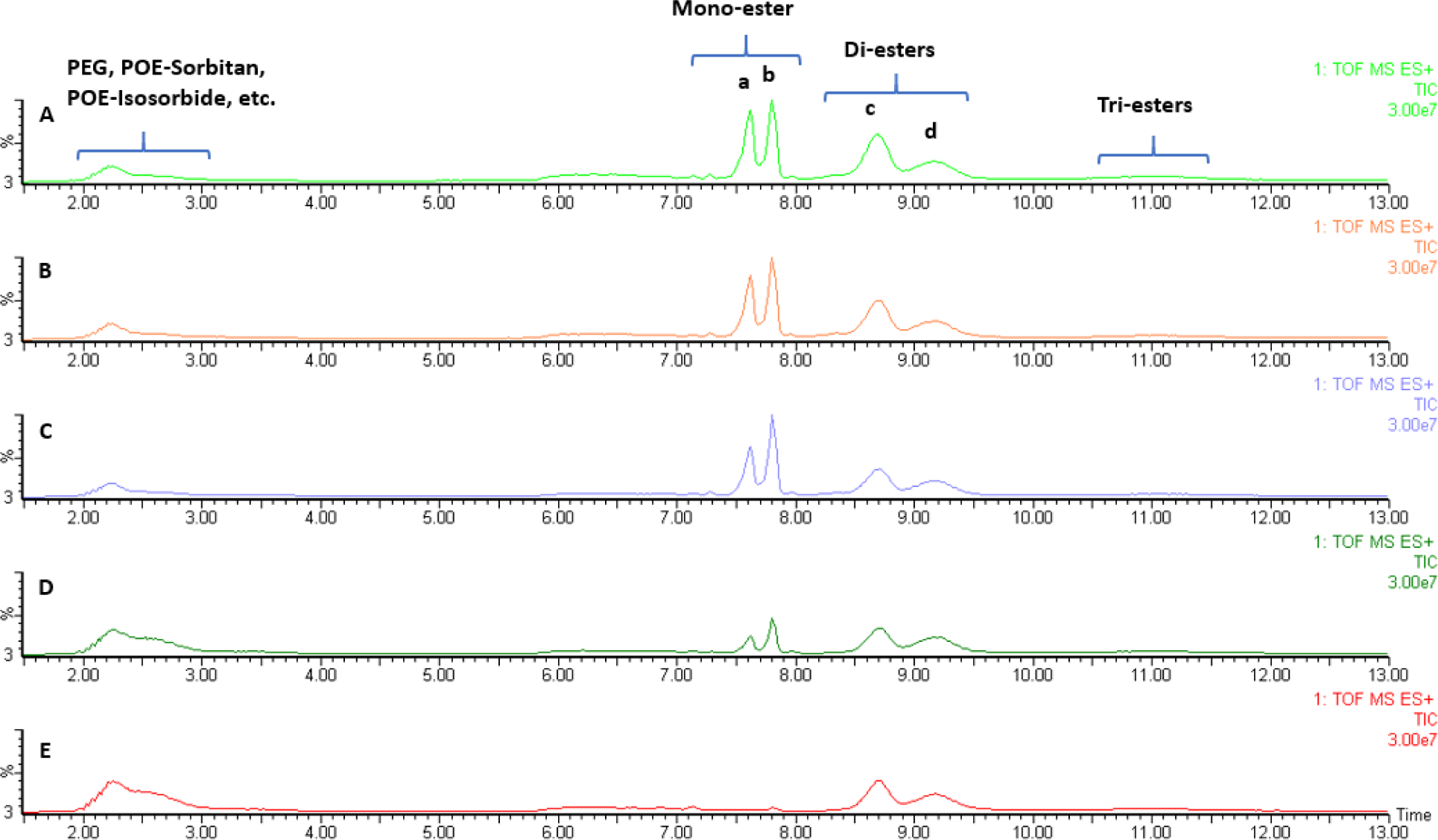
Characterization of PS-80 degradation by LC-MS. Various recombinant enzymes (10 ppm) were added in mAb2 DS from an improved process and incubated with 0.02% PS-80. PS-80 purity were measured by RP-UHPLC-ESI(+)-HRMS on day 7 from samples with or without spike-in enzyme. Mono-esters from PS-80 were the major species degraded by LPLA2 and PLA2G7. A. Without recombinant enzyme, B. PLBL2, C. SIAE, D. PLA2G7, E. LPLA2.

## Discussion

Serine hydrolases are a large functionally diverse group of enzymes with great biological and pharmaceutical importance.^21, 30^ By using an activated serine as a nucleophile, these enzymes hydrolyze ester, thioester and amide bonds in a variety of substrates including metabolites, lipids, peptides and proteins. For this reason, they play important roles in physiological and pathological systems.^21, 30^ The serine hydrolase superfamily is subdivided among lipases, esterases, thioesterases, amidases and peptidases.^21, 30^ In pharmaceutical bioprocess, serine hydrolases secreted from host cell lines or released from broken cells during cell culture have been reported to significantly impact the biological drug substance and its corresponding formulation excipients. For example, complement component 1’s (C1s), a serine protease from the complement cascade, was shown to proteolyze HIV surface protein gp120 in CHO cells,^33^ which have hindered the development of biologics including HIV vaccines.^33^ Several serine hydrolases, including LPLA2, LPL, have been reported to degrade the formulation excipient PS.^5-7^ Problematic residual proteases and lipases are usually identified by TABP, the sensitivity of which is limited by the high abundance of the drug substance in a sample after extensive purification. However, even low concentrations of the HCP contaminants can be problematic. The effect of contaminant proteins was demonstrated by spiking partially purified recombinant proteins at higher concentrations than typically detected in endogenous samples. Moreover, not all proteases and lipases have the same activity at the same concentration. It is possible that the most active hydrolases can cause noticeable damage while being below the detection limit of the TABP assay. Hence, it is critical to develop a method that can enrich problematic classes of proteins for process characterization and root cause investigation.

Based on the mechanism of action of serine hydrolases (Figure 1), ABPP has been developed to specifically enrich and identify proteins from this superfamily.^21, 30^ Covalent protein modifiers play a key role as starting points for designing irreversible enzyme inhibitors and developing chemical probes for ABPP.^26^ To optimize ABPP for serine hydrolases profiling in HCCF from biologics cell culture, two previously reported FP probes were evaluated using the same workflow. As shown in Figure 2, FP-Biotin probe has better enrichment for most serine hydrolases compared with FP-Desthiobiotin probe. The commercial FP-Desthiobiotin probe was used in the recent application of ABPP for PS-80 degradation.^29^ It is possible that different chemical probes have different affinity for various serine hydrolyses. There has been significant research focused on chemical probe development for selected groups of serine hydrolyses or for different applications.^34-35^ Theoretically, various chemical probes with different reactive groups but the same purification group may be used to widen the detection range of serine hydrolases.^23^ In this study the variation in enrichment is most likely due to the differential affinity of biotin and desthiobiotin for streptavidin. Not limited to serine hydrolases, ABPP assays can be used to identify active enzymes from other classes including cysteine-,threonine-, aspartyl- or metalloproteases and glycosidases using similar workflow but using different chemical probes.^22, 36^ Those enzymes may have similar physicochemical properties as some biotherapeutic proteins, or bind to biotherapeutics for co-purification. For example, cathepsin D, a member of the aspartyl protease family, caused antibody fragment or particle formation in monoclonal antibody products.^37-38^ In another example, cathepsin L, a cysteine protease, caused proteolytic cleavage of CHO expressed proteins during processing and storage.^39^ In those studies, extensive enrichment was performed in order to identify those low abundance HCP enzymes.^37-39^ There is opportunity to develop a universal ABPP approach to identify a dozen members of the cathepsin family using a pool of chemical probes for active serine, cysteine and aspartyl proteases.

To the best of our knowledge, this study demonstrated the first application of an optimized ABPP approach for serine hydrolase profiling in cell culture fluid from cells used to produce therapeutic proteins. The composition and activity of serine hydrolases will help understand the source of those HCPs and their impact on the stability of the biologics drug substance and its formulation excipients. ABPP could also help identify and annotate proteins with potential new functions depending on the specific chemical probes used. The understanding of serine hydrolases, especially those known to have adverse impacts, such as proteases and lipases (Figure 5), will provide useful information for cell line development, clone selection, as well as upstream and downstream development. In our study, the serine hydrolase HCPs from two mAbs showed distinct profiles (Figure 6). The difference may result from different strains of CHO cell lines, from differences in the cell culture conditions, or from downstream purification steps. This may provide a rationale to evaluate serine hydrolase and lipase composition for each therapeutic protein.

In this study, ABPP is also used to investigate PS degradation observed in biologics formulation. There is an industry-wide challenge to identify trace levels of lipases/esterase causing PS-80 degradation. Those active enzymes usually fall below the limits of detection of TABP assay in drug substance.^6^ Also, assessment of protein abundance may not correlate with the amount of active enzyme present in a sample showing PS degradation. Currently, there is no direct, high-throughput assay to assess lipase activity in biotherapeutics. For example, our current assay for measuring PS-80 degradation requires 1-3 weeks, includes incubating samples at 37 °C and monitoring degradation by HPLC or LC-MS. ABPP potentially addresses both abundance and activity challenges by identifying those low abundant enzymes based on their activity rather than abundance.

As a proof-of-concept study, process intermediates from two mAbs were tested by both TABP and ABPP analysis. PLBL2 was the only lipase identified by TABP in mAb1 as shown in Table 3, but it did not appear to cause PS-80 degradation. PLBL2 was also not identified as an active lipase in HCCF samples from mAb1 or mAb2 (Figure 2). Recombinant CHO PLBL2 at concentrations up to 10 ppm was confirmed to have no activity on PS-80 degradation as shown in Figure 7. Early study has suggested PLBL2 to be the root cause of PS-20 degradation in a sulfatase drug product;^4^ however, the amount of PLBL2 (90% pure from a non-CHO species) used for functional confirmation was much higher than the concentration observed in the drug product sample.^4^ The lack of degradation seen in our study in the presence of PLBL2 contaminants is consistent with early reports that purified recombinant CHO PLBL2 or mAb biotherapeutic in-process samples showed no *in vitro* phospholipase activity against synthetic substrates.^13^ Actually, bovine phospholipase B□like protein was proposed to be an amidase or peptidase rather than a lipase. No natural substrate for the members of the PLBD family has been yet reported.^40^ Our data suggests that CHO PLBL2 is neither an amidase nor a peptidase in the serine hydrolase family due to the lack of evidence of this activity in our data. There are still possibilities that PLBL2 present in the sample in an inactive or inhibited conformation that is not enriched by the fluorophosphate chemical probe. Overall, the impact of PLBL2 on PS degradation at concentrations detected in biotherapeutics is questionable according to our data, although the level of PLBL2 must be controlled due to an immunogenicity risk.^13^ This observation is consistent with recent report that PLBL2 is not responsible for PS degradation in mAb drug products.^41^ We have observed protein formulations containing over 500 ppm of PLBL2 without any PS-80 degradation (unpublished results). The role of different pH, buffer conditions, post-transitional modifications and association with drug substance for PLBL2 on PS degradation may need further study.

In contrast, there was significant PS-80 degradation observed in mAb2, but no lipase or esterase were identified by TABP while ABPP identified two lipases/esterases. The first lipase/esterase detected, PLA2G7, a member of the phospholipase A2 family also known as platelet-activating factor (PAF) acetylhydrolase, modulates the action of PAF by hydrolyzing the sn-2 ester bond to yield the biologically inactive lyso-PAF.^42^ Another member of the phospholipase A2 family, LPLA2 (PLA2G15), has been reported to degrade both PS-80 and PS-20 at concentrations as low as 0.3 ppm.^6^ Both lipases have been confirmed to be able to degrade PS-80 in mAb2 (Figure 7). The second esterase identified by ABPP from mAb2, SIAE (sialic acid acetylesterase) belongs to the family of hydrolases, specifically acting on carboxylic ester bonds, which is the same ester bond as in PS-80.^43^ The *in vitro* incubation study demonstrated limited activity for recombinant SIAE against PS-80 degradation (Figure 7). The lack of activity may due to the quality or selectivity of the recombinant enzyme, which needs to be further investigated using high-quality recombinant active CHO enzymes, specific inhibitors or genetic knockout cell lines. The successful identification of novel lipases for PS-80 degradation by ABPP demonstrates the potential of ABPP for root cause investigation of PS degradation in bioprocess samples.

The ability of ABPP to identify low-abundance problematic enzymes responsible for PS degradation may lead to the development of specific enzymatic and/or ELISA assays to measure and control the enzyme levels. It may also help develop novel processes, formulations, or genetic strategies to remove or de-activate those enzymes in final drug product to ensure a sufficiently safe final drug product. The ability to confirm if certain lipases, such as PLBL2, are active may lead to modification of the acceptable levels of some problematic HCPs for product development, which may significantly reduce the associated cost of manufacturing and place a decreased burden on process validation.

## Conclusions

In summary, there is significant analytical challenge to identify low abundant active enzymes responsible for PS degradation in biotherapeutics formulations. Based on the mechanism of action of serine hydrolases, we have established an optimized ABPP approach for profiling active serine hydrolases from bioprocess samples using active chemical probes. ABPP benefits from monitoring and enriching the active forms of enzymes and providing greater sensitivity over TABP for identifying active residual host cell enzymes of lower abundance. Our ABPP assay did not enrich PLBL2, suggesting that it might not be a serine hydrolase or may not be active, and thus not contributing to PS-80 degradation. From the proof-of-concept study for PS-80 degradation, PLA2G7 was identified and confirmed as a novel lipase for PS 80 degradation by ABPP. Taken together, ABPP expands the existing analytical toolbox for HCP characterization, and demonstrates its unique potential to advance biologics process and product development.

## Acknowledgments

The authors are grateful to Li Zheng and Adam Schwaid for discussion during method development, and Shannon Rivera for reviewing and feedback of the manuscript. We thank John Welsh, Rebecca Chmielowski and Justin Miller for sharing the recombinant PLBL2 and LPLA2 enzymes.

## Competing Interest Statement

All authors are employees of Merck Sharp & Dohme Corp., a subsidiary of Merck & Co., Inc., Kenilworth, NJ, USA. XL, SL, DDR, GCA, CA and SB are listed inventors on U.S. Provisional Application Serial No. 63/043,966, Filed on June 25, 2020 entitled “SERINE HYDROLASE PROFILING ASSAY IN BIOTHEURAPEUTIC”.

## References

1. Dwivedi, M., Blech, M., Presser, I., Garidel, P., Polysorbate degradation in biotherapeutic formulations: Identification and discussion of current root causes. Int J Pharm 2018, 552 (1-2), 422–436.

2. Kerwin, B. A., Polysorbates 20 and 80 used in the formulation of protein biotherapeutics: structure and degradation pathways. J Pharm Sci2008, 97 (8), 2924–35.

3. Kishore, R. S., Kiese, S., Fischer, S., Pappenberger, A., Grauschopf, U., Mahler, H. C., The degradation of polysorbates 20 and 80 and its potential impact on the stability of biotherapeutics. Pharm Res2011, 28 (5), 1194–210.

4. Dixit, N., Salamat-Miller, N., Salinas, P. A., Taylor, K. D., Basu, S. K., Residual Host Cell Protein Promotes Polysorbate 20 Degradation in a Sulfatase Drug Product Leading to Free Fatty Acid Particles. J Pharm Sci2016, 105 (5), 1657–1666.

5. Chiu, J., Valente, K. N., Levy, N. E., Min, L., Lenhoff, A. M., Lee, K. H., Knockout of a difficult-to-remove CHO host cell protein, lipoprotein lipase, for improved polysorbate stability in monoclonal antibody formulations. Biotechnol Bioeng2017, 114 (5), 1006–1015.

6. Hall, T., Sandefur, S. L., Frye, C. C., Tuley, T. L., Huang, L., Polysorbates 20 and 80 Degradation by Group XV Lysosomal Phospholipase A2 Isomer X1 in Monoclonal Antibody Formulations. J Pharm Sci2016, 105 (5), 1633–1642.

7. McShan, A. C., Kei, P., Ji, J. A., Kim, D. C., Wang, Y. J., Hydrolysis of Polysorbate 20 and 80 by a Range of Carboxylester Hydrolases. PDA J Pharm Sci Technol2016, 70 (4), 332–45.

8. Tomlinson, A., Demeule, B., Lin, B., Yadav, S., Polysorbate 20 Degradation in Biopharmaceutical Formulations: Quantification of Free Fatty Acids, Characterization of Particulates, and Insights into the Degradation Mechanism. Mol Pharm2015, 12 (11), 3805–15.

9. Labrenz, S. R., Ester hydrolysis of polysorbate 80 in mAb drug product: evidence in support of the hypothesized risk after the observation of visible particulate in mAb formulations. J Pharm Sci2014, 103 (8), 2268–77.

10. Tran, B., Grosskopf, V., Wang, X., Yang, J., Walker, D., Jr., Yu, C., McDonald, P., Investigating interactions between phospholipase B-Like 2 and antibodies during Protein A chromatography. J Chromatogr A 2016, 1438, 31–8.

11. Bracewell, D. G., Francis, R., Smales, C. M., The future of host cell protein (HCP) identification during process development and manufacturing linked to a risk-based management for their control. Biotechnol Bioeng2015, 112 (9), 1727–37.

12. Gao, X., Rawal, B., Wang, Y., Li, X., Wylie, D., Liu, Y. H., Breunig, L., Driscoll, D., Wang, F., Richardson, D. D., Targeted Host Cell Protein Quantification by LC-MRM Enables Biologics Processing and Product Characterization. Anal Chem2020, 92 (1), 1007–1015.

13. Fischer, S. K., Cheu, M., Peng, K., Lowe, J., Araujo, J., Murray, E., McClintock, D., Matthews, J., Siguenza, P., Song, A., Specific Immune Response to Phospholipase B-Like 2 Protein, a Host Cell Impurity in Lebrikizumab Clinical Material. AAPS J2017, 19 (1), 254–263.

14. Vanderlaan, M., Zhu-Shimoni, J., Lin, S., Gunawan, F., Waerner, T., Van Cott, K. E., Experience with host cell protein impurities in biopharmaceuticals. Biotechnol Prog2018, 34 (4), 828–837.

15. Walker, D. E., Yang, F., Carver, J., Joe, K., Michels, D. A., Yu, X. C., A modular and adaptive mass spectrometry-based platform for support of bioprocess development toward optimal host cell protein clearance. MAbs2017, 9 (4), 654–663.

16. Kufer, R., Haindl, M., Wegele, H., Wohlrab, S., Evaluation of Peptide Fractionation and Native Digestion as Two Novel Sample Preparation Workflows to Improve HCP Characterization by LC-MS/MS. Anal Chem2019, 91 (15), 9716–9723.

17. Huang, L., Wang, N., Mitchell, C. E., Brownlee, T., Maple, S. R., De Felippis, M. R., A Novel Sample Preparation for Shotgun Proteomics Characterization of HCPs in Antibodies. Anal Chem2017, 89 (10), 5436–5444.

18. Yang, F., Walker, D. E., Schoenfelder, J., Carver, J., Zhang, A., Li, D., Harris, R., Stults, J. T., Yu, X. C., Michels, D. A., A 2D LC-MS/MS Strategy for Reliable Detection of 10-ppm Level Residual Host Cell Proteins in Therapeutic Antibodies. Anal Chem2018, 90 (22), 13365–13372.

19. Chen, I. H., Xiao, H., Daly, T., Li, N., Improved Host Cell Protein Analysis in Monoclonal Antibody Products through Molecular Weight Cutoff Enrichment. Anal Chem2020, 92 (5), 3751–3757.

20. Adam, G. C., Sorensen, E. J., Cravatt, B. F., Chemical strategies for functional proteomics. Mol Cell Proteomics2002, 1 (10), 781–90.

21. Simon, G. M., Cravatt, B. F., Activity-based proteomics of enzyme superfamilies: serine hydrolases as a case study. J Biol Chem2010, 285 (15), 11051–5.

22. Sanman, L. E., Bogyo, M., Activity-based profiling of proteases. Annu Rev Biochem 2014, 83, 249–73.

23. Adam, G. C., Sorensen, E. J., Cravatt, B. F., Proteomic profiling of mechanistically distinct enzyme classes using a common chemotype. Nat Biotechnol2002, 20 (8), 805–9.

24. Liu, Y., Patricelli, M. P., Cravatt, B. F., Activity-based protein profiling: the serine hydrolases. Proc Natl Acad Sci U S A1999, 96 (26), 14694–9.

25. Greenbaum, D., Medzihradszky, K. F., Burlingame, A., Bogyo, M., Epoxide electrophiles as activity-dependent cysteine protease profiling and discovery tools. Chem Biol2000, 7 (8), 569-81. 26.

26. Cravatt, B. F., Wright, A. T., Kozarich, J. W., Activity-based protein profiling: from enzyme chemistry to proteomic chemistry. Annu Rev Biochem 2008, 77, 383–414.

27. Zweerink, S., Kallnik, V., Ninck, S., Nickel, S., Verheyen, J., Blum, M., Wagner, A., Feldmann, I., Sickmann, A., Albers, S. V., Brasen, C., Kaschani, F., Siebers, B., Kaiser, M., Activity-based protein profiling as a robust method for enzyme identification and screening in extremophilic Archaea. Nat Commun 2017, 8, 15352.

28. Lentz, C. S., Sheldon, J. R., Crawford, L. A., Cooper, R., Garland, M., Amieva, M. R., Weerapana, E., Skaar, E. P., Bogyo, M., Identification of a S. aureus virulence factor by activity-based protein profiling (ABPP). Nat Chem Biol2018, 14 (6), 609–617.

29. Zhang, S., Xiao, H., Molden, R., Qiu, H., Li, N., Rapid Polysorbate 80 Degradation by Liver Carboxylesterase in a Monoclonal Antibody Formulated Drug Substance at Early Stage Development. J Pharm Sci 2020.

30. Long, J. Z., Cravatt, B. F., The metabolic serine hydrolases and their functions in mammalian physiology and disease. Chem Rev2011, 111 (10), 6022–63.

31. Adibekian, A., Martin, B. R., Chang, J. W., Hsu, K. L., Tsuboi, K., Bachovchin, D. A., Speers, A. E., Brown, S. J., Spicer, T., Fernandez-Vega, V., Ferguson, J., Hodder, P. S., Rosen, H., Cravatt, B. F., Confirming target engagement for reversible inhibitors in vivo by kinetically tuned activity-based probes. J Am Chem Soc2012, 134 (25), 10345–8.

32. Cheng, Y., Hu, M., Zamiri, C., Carcelen, T., Demeule, B., Tomlinson, A., Gu, J., Yigzaw, Y., Kalo, M., Yu, X. C., A Rapid High-Sensitivity Reversed-Phase Ultra High Performance Liquid Chromatography Mass Spectrometry Method for Assessing Polysorbate 20 Degradation in Protein Therapeutics. J Pharm Sci2019, 108 (9), 2880–2886.

33. Li, S. W., Yu, B., Byrne, G., Wright, M., O’Rourke, S., Mesa, K., Berman, P. W., Identification and CRISPR/Cas9 Inactivation of the C1s Protease Responsible for Proteolysis of Recombinant Proteins Produced in CHO Cells. Biotechnol Bioeng2019, 116 (9), 2130–2145.

34. Wang, C., Abegg, D., Dwyer, B. G., Adibekian, A., Discovery and Evaluation of New Activity-Based Probes for Serine Hydrolases. Chembiochem2019, 20 (17), 2212–2216.

35. Adam, G. C., Cravatt, B. F., Sorensen, E. J., Profiling the specific reactivity of the proteome with non-directed activity-based probes. Chem Biol2001, 8 (1), 81–95.

36. Yang, P., Liu, K., Activity-based protein profiling: recent advances in probe development and applications. Chembiochem2015, 16 (5), 712–24.

37. Bee, J. S., Tie, L., Johnson, D., Dimitrova, M. N., Jusino, K. C., Afdahl, C. D., Trace levels of the CHO host cell protease cathepsin D caused particle formation in a monoclonal antibody product. Biotechnol Prog2015, 31 (5), 1360–9.

38. Lim, A., Doyle, B. L., Kelly, G. M., Reed-Bogan, A. M., Breen, L. H., Shamlou, P. A., Lambooy, P. K., Characterization of a cathepsin D protease from CHO cell-free medium and mitigation of its impact on the stability of a recombinant therapeutic protein. Biotechnol Prog2018, 34 (1), 120–129.

39. Luo, H., Tie, L., Cao, M., Hunter, A. K., Pabst, T. M., Du, J., Field, R., Li, Y., Wang, W. K., Cathepsin L Causes Proteolytic Cleavage of Chinese-Hamster-Ovary Cell Expressed Proteins During Processing and Storage: Identification, Characterization, and Mitigation. Biotechnol Prog2019, 35 (1), e2732.

40. Repo, H., Kuokkanen, E., Oksanen, E., Goldman, A., Heikinheimo, P., Is the bovine lysosomal phospholipase B-like protein an amidase? Proteins2014, 82 (2), 300–11.

41. Zhang, S., Xiao, H., Goren, M., Burakov, D., Chen, G., Li, N., Tustian, A., Adams, B., Mattila, J., Bak, H., Putative Phospholipase B-Like 2 is Not Responsible for Polysorbate Degradation in Monoclonal Antibody Drug Products. J Pharm Sci2020, 109 (9), 2710–2718.

42. Tjoelker, L. W., Eberhardt, C., Unger, J., Trong, H. L., Zimmerman, G. A., McIntyre, T. M., Stafforini, D. M., Prescott, S. M., Gray, P. W., Plasma platelet-activating factor acetylhydrolase is a secreted phospholipase A2 with a catalytic triad. J Biol Chem1995, 270 (43), 25481–7.

43. Schauer, R., Reuter, G., Stoll, S., Sialate O-acetylesterases: key enzymes in sialic acid catabolism. Biochimie1988, 70 (11), 1511–9.

